# Loss of IL-10 signaling promotes IL-22 dependent host defenses against acute *Clostridioides difficile* infection

**DOI:** 10.1101/2020.10.14.340190

**Authors:** Emily S. Cribas, Joshua E. Denny, Jeffrey R. Maslanka, Michael C. Abt

**Affiliations:** Department of Microbiology, Perelman School of Medicine, University of Pennsylvania, Philadelphia, Pennsylvania, USA

## Abstract

Infection with the bacterial pathogen *Clostridioides difficile* causes severe damage to the intestinal epithelium that elicits a robust inflammatory response. Markers of intestinal inflammation accurately predict clinical disease severity. However, determining the extent to which host-derived proinflammatory mediators drive pathogenesis versus promote host protective mechanisms remains elusive. In this report, we employed *Il10^-/-^* mice as a model of spontaneous colitis to examine the impact of constitutive intestinal immune activation, independent of infection, on *C. difficile* disease pathogenesis. Upon *C. difficile* challenge, *Il10^-/-^* mice exhibited significantly decreased morbidity and mortality compared to littermate *Il10* heterozygote (*Il10*^HET^) control mice, despite a comparable *C. difficile* burden, innate immune response, and microbiota composition following infection. Similarly, antibody-mediated blockade of IL-10 signaling in wild-type C57BL/6 mice conveyed a survival advantage if initiated three weeks prior to infection. In contrast, no advantage was observed if blockade was initiated on the day of infection, suggesting that constitutive activation of inflammatory defense pathways prior to infection mediated host protection. IL-22, a cytokine critical in mounting a protective response against *C. difficile* infection, was elevated in the intestine of uninfected, antibiotic-treated *Il10^-/-^* mice, and genetic ablation of the IL-22 signaling pathway in *Il10^-/-^* mice negated the survival advantage following *C. difficile* challenge. Collectively, these data demonstrate that constitutive loss of IL-10 signaling, via genetic ablation or antibody blockade, enhances IL-22 dependent host defense mechanisms to limit *C. difficile* pathogenesis.

## Introduction

*Clostridioides difficile* is a leading cause of nosocomial infections in the United States. High recurrence rates, increases in community-acquired infections, and the emergence of antibiotic-resistant strains render *C. difficile* an urgent threat to our public health system^1–5^. The manifestation of *C. difficile* infection is highly variable; ranging from asymptomatic colonization, diarrhea, and pseudomembranous colitis, to severe cases of toxic megacolon and death^6^. Disease severity is shaped by the host immune response and patients on immunosuppressants or with autoimmune disorders are more susceptible to severe disease^7–9^. Taken together, there is a need to study the host immune response to *C. difficile* infection to develop new therapies.

Upon intestinal colonization, *C. difficile* produces toxins that disrupt epithelial barrier integrity and result in translocation of commensal bacteria into submucosal tissues. Impaired barrier integrity leads to downstream induction of a multi-faceted, robust, inflammatory response^10,11^. The innate immune response is essential for protection against *C. difficile* infection. Mice deficient in pathogen recognition receptor signaling pathways or innate immune cells exhibit increased bacterial translocation, damage to the epithelial barrier, and increased mortality following *C. difficile* infection^12–16^. Conversely, proinflammatory mediators can simultaneously exacerbate tissue damage and promote *C. difficile* expansion to hinder recovery^16–18^. In support of these animal studies, elevated fecal and serum proinflammatory cytokine levels are associated with increased disease severity in patients^19–21^. Together, these findings highlight the complexity of the host response and demonstrate the need to fundamentally understand the timing and context of intestinal inflammation as a driver of *C. difficile* pathogenesis. To begin to address the contribution of the host proinflammatory immune response in promoting disease severity during *C. difficile* infection, elevated expression of intestinal inflammatory mediators was established in mice *a priori C. difficile* challenge and the disease severity following subsequent infection was investigated.

Interleukin-10 (IL-10) is a broad immunoregulatory cytokine that negatively regulates commensal bacteria-driven immune activation at steady-state^22–24^. Intestinal expression of IL-10 is critical for maintaining intestinal homeostasis as mice deficient in the *Il10* gene develop microbiota-dependent spontaneous colitis characterized by chronic activation of inflammatory mediators that are also associated with *C. difficile* pathogenesis^25–27^. Thus, *Il10*^-/-^ mice, a widely used model of intestinal immune dysregulation, offer the opportunity to decouple intestinal inflammation from infection to study the causative nature of inflammatory mediators in *C. difficile* pathogenesis^28^.

In this report, we demonstrate that pre-existing intestinal immune activation, e.g. expression of proinflammatory cytokines driven by loss of IL-10 signaling, reduces susceptibility to *C. difficile* infection. Host protective immunity was independent of changes in *C. difficile* burden, toxin production, or the microbiota. The protective capacity of IL-10-deficient immune activation was dependent on IL-22 production enhancing early host defenses against *C. difficile* infection.

## Results

### IL-10 deficiency decreases susceptibility to acute *C. difficile* infection

At steady-state, IL-10 maintains intestinal homeostasis by negatively regulating commensal bacteria-driven expression of proinflammatory cytokines. In the context of *C. difficile* infection, many of these proinflammatory cytokines correlate with increased disease severity. However, it is unclear whether the inflammatory profile associated with infection emerges following *C. difficile*-mediated tissue damage or if it proactively drives pathology and worsens disease^20,21^. To address this question, intestinal inflammation was induced independently of *C. difficile* infection using the murine *Il10^-/-^* spontaneous colitis model and the impact of constitutive inflammation on *C. difficile* disease severity was examined. Cohoused *Il10^-/-^* and littermate *Il10* heterozygous mice (*Il10*^HET^) were treated with a broad-spectrum antibiotic cocktail in their drinking water to induce susceptibility to *C. difficile* and mimic the microbiota dysbiosis observed in patients at high risk for contracting *C. difficile.* antibiotic-treated *Il10^HET^* mice, exhibited peak disease severity within 48 hours of infection as measured by a disease score that measures weight loss, body temperature, diarrhea, and lethargy (Fig. 1A), and approximately 75% mortality rate (Fig. 1B). In contrast, *Il10^-/-^* mice experienced reduced disease severity at 2 days post infection (p.i.) (Fig. 1A) and were less likely to succumb to acute *C. difficile* infection compared to *Il10^HET^* mice (Fig. 1B).

*Il10^-/-^* mice challenged with pathogenic *Escherichia coli, Salmonella typhimurium, Citrobacter rodentium, Toxoplasma gondii*, or *Candida albicans* all display improved pathogen clearance via enhanced phagocytic mechanisms by innate immune cells^29–33^. Thus, *C. difficile* burden was measured at day 1 and 2 p.i. No difference in *C. difficile* burden was observed in the cecal content of *Il10*^HET^ and *Il10^-/-^* mice at days 1 (Fig. 1C) or 2 p.i. (Fig. 1D). Further, *C. difficile* toxin activity in the cecal content of *Il10*^HET^ and *Il10^-/-^* mice was similar at day 2 p.i. as measured by an *in vitro* cell rounding assay (Fig. 1E). Together, these data indicate that loss of IL-10 augments host immunity following *C. difficile* infection but does not alter establishment of infection or production of toxins, the primary virulence factors of *C. difficile*.

**Figure 1:**
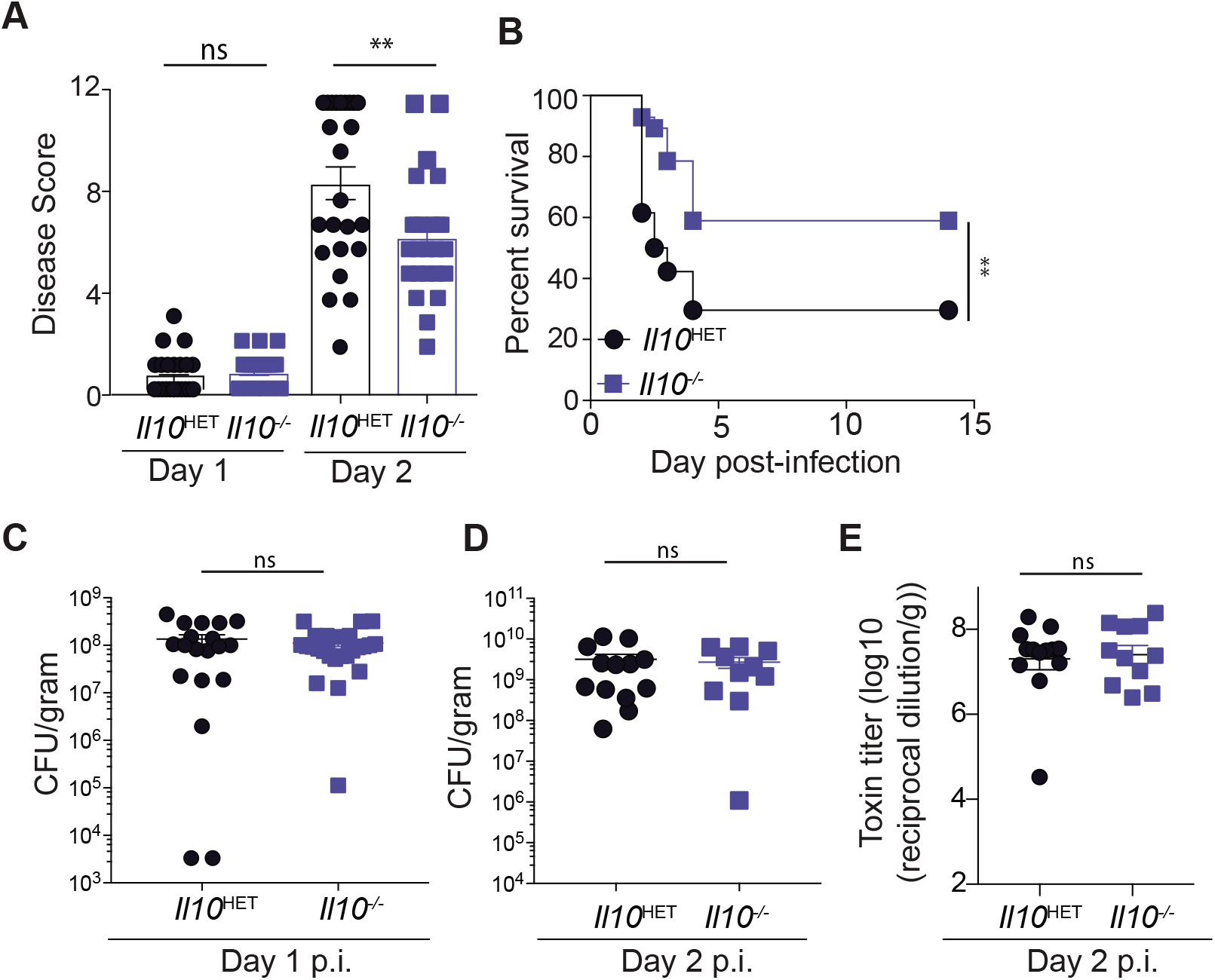
Genetic ablation of *Il10* results in reduced susceptibility to *C. difficile* infection. Antibiotic-treated *Il10^-/-^* and *Il10*^HET^ mice were inoculated with approximately 400 spores of *C. difficile* (VPI 10463 strain) and monitored daily for disease. **(A)** Disease score and **(B)** survival following infection. Data shown are a combination of five independent experiments (*Il10*^-/-^,n=23; *Il10*^HET^, n=25). **(C)** *C. difficile* burden in fecal pellets at day 1 p.i. **(D)** *C. difficile* burden and **(E)** *C. difficile* toxin levels in the cecal content at day 2 p.i. ** = p<0.01. Statistical significance was calculated by a log-rank test.

Intestinal inflammation and subsequent onset of spontaneous colitis in *Il10^-/-^* mice varies between vivaria and is dependent on the microbiota^25,26^. To test the rigor of the observed phenotype in *C. difficile* infected *Il10^-/-^* mice, complimentary experiments with cohoused wild-type C57BL/6 and *Il10*^-/-^ mice were conducted in an independent animal facility. In agreement with our studies in *Il10*^HET^ mice, *Il10^-/-^* mice exhibited improved survival (Suppl. Fig. 1A) compared to cohoused C57BL/6 mice following *C. difficile* infection despite no difference in *C. difficile* burden (Suppl. Fig. 1B) or toxin production (Suppl. Fig. 1C), demonstrating the robustness of this phenotype.

### Enhanced protection in *Il10^-/-^* mice is not driven by a distinct microbiota composition

The composition of the microbiota impacts *C. difficile* pathogenesis through multiple direct and indirect mechanisms^34^, therefore cohoused littermate mice were used to normalize for this variable. To test the null hypothesis that the microbiota composition between *Il10*^HET^ and *Il10^-/-^* mice was indistinguishable, bacterial 16S rRNA marker gene profiling was conducted on cecal content from *Il10*^HET^ and *Il10^-/-^* mice collected at day 2 post *C. difficile* or mock infection. Microbial community alpha diversity was not different between uninfected or infected *Il10*^HET^ and *Il10^-/-^* mice (Fig. 2A). Comparison of 16S rRNA bacterial community profiles between *Il10*^HET^ and *Il10^-/-^* mice by relative bacterial abundance revealed a bloom of amplicon sequence variants (ASVs) identified as *C. difficile* in both *Il10*^HET^ and *Il10^-/-^* infected mice compared to uninfected mice, however the relative abundance composition between infected groups was similar (Fig. 2B). Beta diversity comparisons between samples by unsupervised hierarchical clustering (Fig. 2C), unweighted UniFrac distances (Fig. 2D), or PERMANOVA analysis (Suppl. Table 1) did not lead us to reject the null hypothesis that there was no microbial community level differences between *Il10*^HET^ and *Il10^-/-^* mice at day 2 following mock infection or *C. difficile* infection. A linear regression model was used to identify individual ASVs that correlate with *Il10*^HET^ and *Il10^-/-^* phenotypes. The linear model readily detected *C. difficile* as significantly enriched in infected mice compared to uninfected mice but failed to identify an ASV significantly different between the microbiota of infected *Il10*^HET^ and *Il10^-/-^* mice (Suppl. Fig. 2A,B).

**Figure 2:**
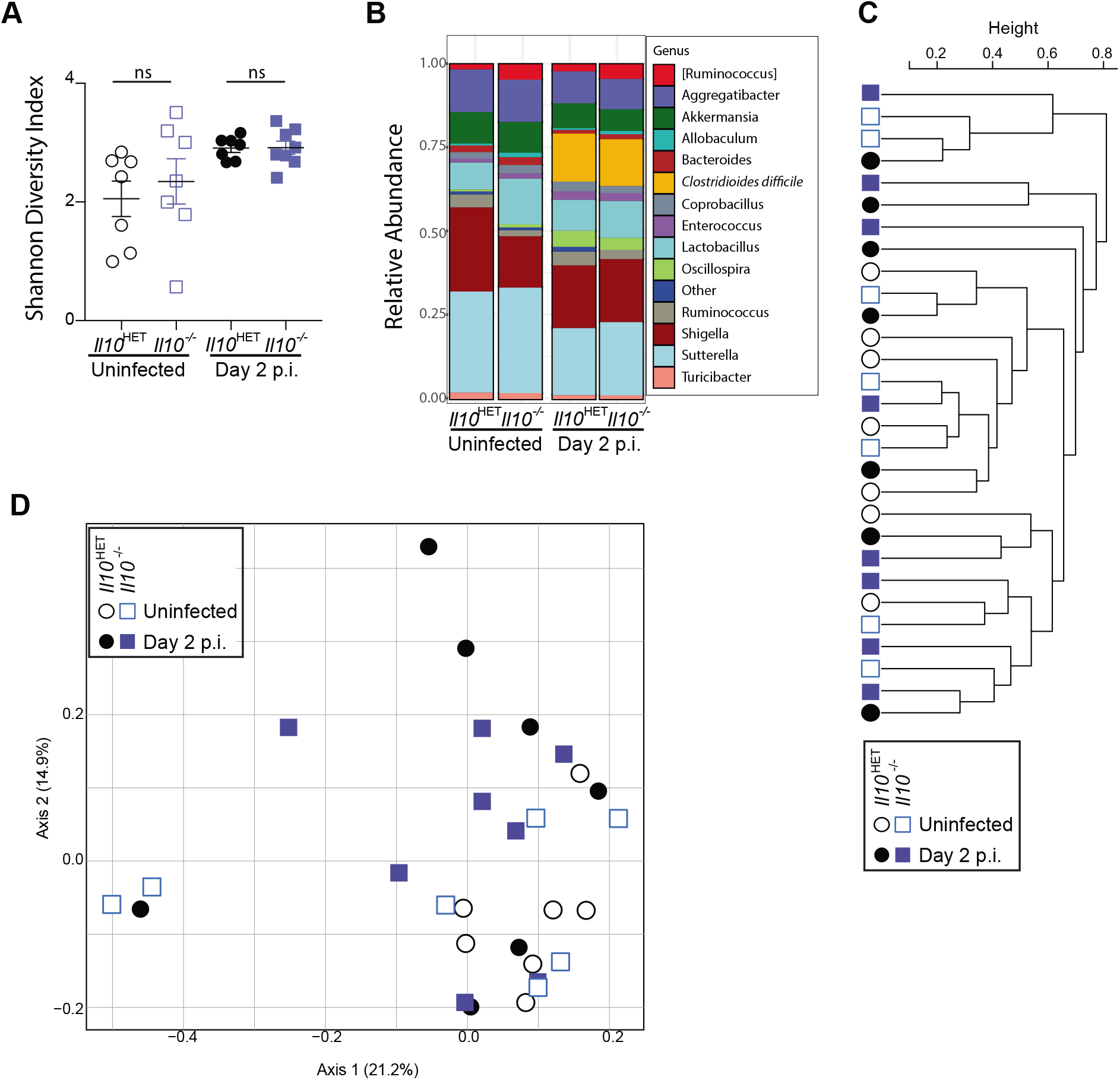
*C. difficile-infected Il10^-/-^* mice and *Il10*^HET^ mice exhibit a similar microbiota composition. Antibiotic-treated uninfected and *C. difficile* infected *Il10^-/-^* mice and *Il10*^HET^ mice were sacrificed at day 2 p.i. and cecal content was processed for 16s rRNA bacterial gene profiling. **(A)** Microbial alpha diversity as determined by the Shannon diversity index. **(B)** Relative abundance of top 15 bacterial ASVs. Bar plot is displayed at the genus level except for orange bars that represent an ASV aligning to *C. difficile*. **(C)** Dendrogram representation of intestinal microbial communities using unsupervised hierarchical clustering of unweighted UniFrac distances to identify similarities between samples. **(D)** Unweighted UniFrac principal coordinate analysis plot of 16S bacterial rRNA ASVs.

In a validation cohort, 16S rRNA marker gene profiling was conducted on fecal pellets collected from C57BL/6 and *Il10^-/-^* mice prior to cohousing (day −64 p.i.), throughout cohousing, the start of antibiotic treatment (day −6 p.i.), and on the day of infection (day 0 p.i.). Prior to cohousing, *Il10^-/-^* mice exhibit a distinct microbiota (Suppl. Fig. 3A, Suppl. Table 2). Cohousing shifted the microbiota of C57BL/6 mice to resemble the microbiota of *Il10^-/-^* mice as determined by unweighted UniFrac distance analysis (Suppl. Fig. 3A), relative bacterial genera abundance (Suppl. Fig. 3B), unsupervised hierarchical clustering (Suppl. Fig. 3C), and PERMANOVA analysis (Suppl. Table 2). Antibiotic treatment between day −6 and 0 p.i. significantly reduced the alpha diversity (Suppl. Fig. 3D) and shifted the microbiota of both C57BL/6 and *Il10^-/-^* mice, but no difference between groups was observed (Suppl. Table 2). To identify specific ASVs that were differentially abundant between cohoused C57BL/6 and *Il10^-/-^* mice, a LEfSe comparison was conducted. Several ASVs were differentially abundant within the microbiota of C57BL/6 and *Il10^-/-^* mice prior to cohousing (Suppl. Fig. 3E). However, following cohousing and antibiotic treatment, none of these differentially abundant ASVs remained (Suppl. Fig. 3E). Together, these microbial profiling data support the conclusion that the differential outcome observed in antibiotic-treated IL-10 sufficient and deficient hosts following *C. difficile* infection cannot be explained by community level differences in the microbiota.

### *Il10^-/-^* and *Il10*^HET^ mice exhibit comparable induction of innate immunity following *C. difficile* infection

No differences in *C. difficile* colonization, toxin production, or microbiota composition were observed between infected *Il10*^HET^ and *Il10^-/-^* mice to account for enhanced protection in *Il10^-/-^* mice, therefore potential immune-mediated mechanisms were assessed. Induction of IL-10 is an effective strategy employed by some enteric pathogens to dampen the host immune response to infection^29,30,35–37^. *C. difficile-derived* flagellin, surface layer proteins, and toxin (TcdB) can all induce macrophages, monocytes, and dendritic cells to produce IL-10 *in vitro*^38–40^. Indeed, C57BL/6 mice infected with *C. difficile* have elevated IL-10 protein in the cecal tissue at day 2 p.i. (Fig. 3A). The broad immunosuppressive functions of IL-10 include inhibition of granulocyte infiltration into mucosal tissue and limiting expression of type-1 and type-17 cytokines, components of the immune response that promote protective immunity following *C. difficile* infection^41–43 44,45^. First protein levels of lipocalin-2 (LCN-2), an established marker of intestinal inflammation^46^, were measured in the cecal content of antibiotic-treated uninfected and day 2 p.i. *Il10^-/-^* and *Il10*^HET^ mice^46^. LCN-2 levels increased to approximately the same concentration in both groups by day 2 p.i (Fig. 3B). To thoroughly assess the quality of the innate immune response to acute *C. difficile* infection, *Il10^-/-^* and *Il10*^HET^ mice were sacrificed at day 2 p.i. and recruitment of innate immune cells and induction of proinflammatory cytokines were assessed. Both infected *Il10^HET^* and *Il10^-/-^* mice exhibited a robust induction of the innate immune response compared to antibiotic-treated, uninfected, control mice (Fig. 3). However, no differences in the frequency (Suppl Fig. 4A,B) or total numbers of infiltrating neutrophils (Fig. 3C), monocytes (Fig. 3D), or eosinophils (Fig. 3E) were observed. *Il10^-/-^* and *Il10*^HET^ mice at day 2 p.i. exhibited comparable elevated gene expression of *Ifng* and *Il22* (Fig. 3F) as well as downstream host defense genes *Nos2* and *Reg3g* in the colon (Fig. 3G). IFN-γ and IL-22 protein concentrations in cecal tissue homogenates were also comparable (Fig. 3H). Type-2 cytokines (IL-5, IL-13, IL-33) associated with eosinophil activation and protection during *C. difficile* infection ^15,38,39^, were not significantly different in the cecum *Il10^-/-^* and *Il10*^HET^ mice at day 2 p.i. (Fig. 3I). Finally, no difference in proinflammatory cytokines (IL-1β, IL-6, IL-17a, IL-27, GM-CSF) reported to modulate *C. difficile* pathogenesis^16,40–44^, or chemokines (CXCL1, CXCL2, CCL2) involved in neutrophil and monocyte recruitment was observed in the cecal tissue homogenates of *Il10^-/-^* and *Il10*^HET^ mice at day 2 p.i. (Suppl. Fig. 4C-D). Collectively, these data indicate the magnitude or quality of the innate immune response in *Il10^-/-^* mice following *C. difficile* infection is not driving the attenuated disease phenotype.

**Figure 3:**
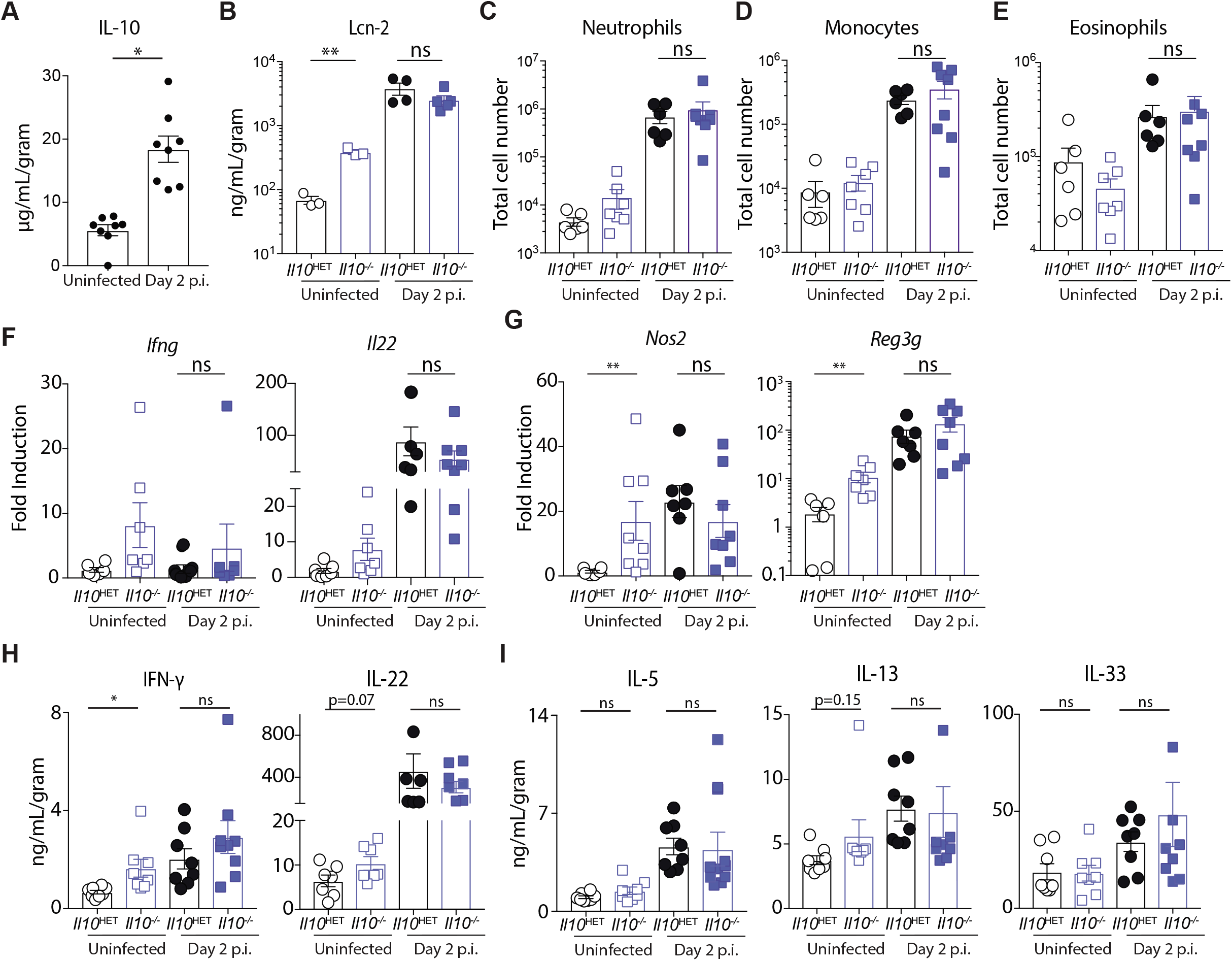
*Il10^-/-^* and *Il10*^HET^ mice exhibit a comparable induction of the innate immune response following acute *C. difficile* infection. **(A)** IL-10 protein levels in the cecal tissue homogenates of antibiotic-treated uninfected and day 2 p.i. C57BL/6 mice. **(B-H)** *Il10^-/-^* and *Il10*^HET^ mice were inoculated with approximately 400 spores of *C. difficile* (VPI 10463 strain) or mock infected and sacrificed two days later. **(B)** LCN-2 protein levels in the cecal supernatants. **(C-E)** Large intestine lamina propria cells were harvested and assessed by flow cytometry for **(C)** neutrophils (CD11b^+^, Ly6G^+^), **(D)** monocyte (CD11b^+^, Ly6C^+^, Ly6G^-^) and **(E)** Eosinophil (SSC^Hi^, CD11b^+^, Siglec-F^+^) recruitment. **(F)** Fold induction of *Ifng, Il22* and **(G)** IFN-γ and IL-22 effector molecules (*Nos2* and *Reg3g*) in the colon at day 2 p.i. relative to uninfected *Il10*^HET^ mice and normalized to *Hprt*. **(H)** IFN-γ, IL-22, and **(I)** type-2 associated cytokine protein levels in the cecal tissue homogenate. Data shown are a combination of two independent experiments (uninfected *Il10*^-/-^,n=7; uninfected *Il10*^HET^, n=6; day 2 infected *Il10*^-/-^,n=8; uninfected *Il10*^HET^, n=7). Data shown are mean ± SEM. * = p<0.05. ** = p<0.01. Statistical significance was calculated by an unpaired t-test.

### Loss of IL-10 signaling prior to *C. difficile* infection drives immune activation in the intestine and augments protective immunity

In contrast to the comparable immune responses observed in *C. difficile* infected *Il10^-/-^* and *Il10*^HET^ mice, antibiotic-treated uninfected *Il10^-/-^* mice at day 2 post mock infection had higher levels of LCN-2 in the cecal content (Fig. 3B.) and increased expression of IL-22 and IFN-γ-dependent effector molecules (Fig. 3F) in the large intestine compared to antibiotic-treated uninfected *Il10*^HET^ mice. Moreover, antibiotic-treated *Il10^-/-^* mice at day 0 p.i. displayed elevated immune activation in the large intestine compared to *Il10*^HET^ mice as determined by increased frequency (Fig. 4A) and total numbers (Fig. 4B) of infiltrating neutrophils in the large intestine as well as elevated expression of proinflammatory immune defense genes (*Il22, Ifng, Reg3g, Nos2*) (Fig. 4C), as has been previously reported^47,48^. These results support the hypothesis that pre-existing immune activation, not the magnitude of the immune response following infection, confers protective immunity in *Il10^-/-^* mice.

**Figure 4:**
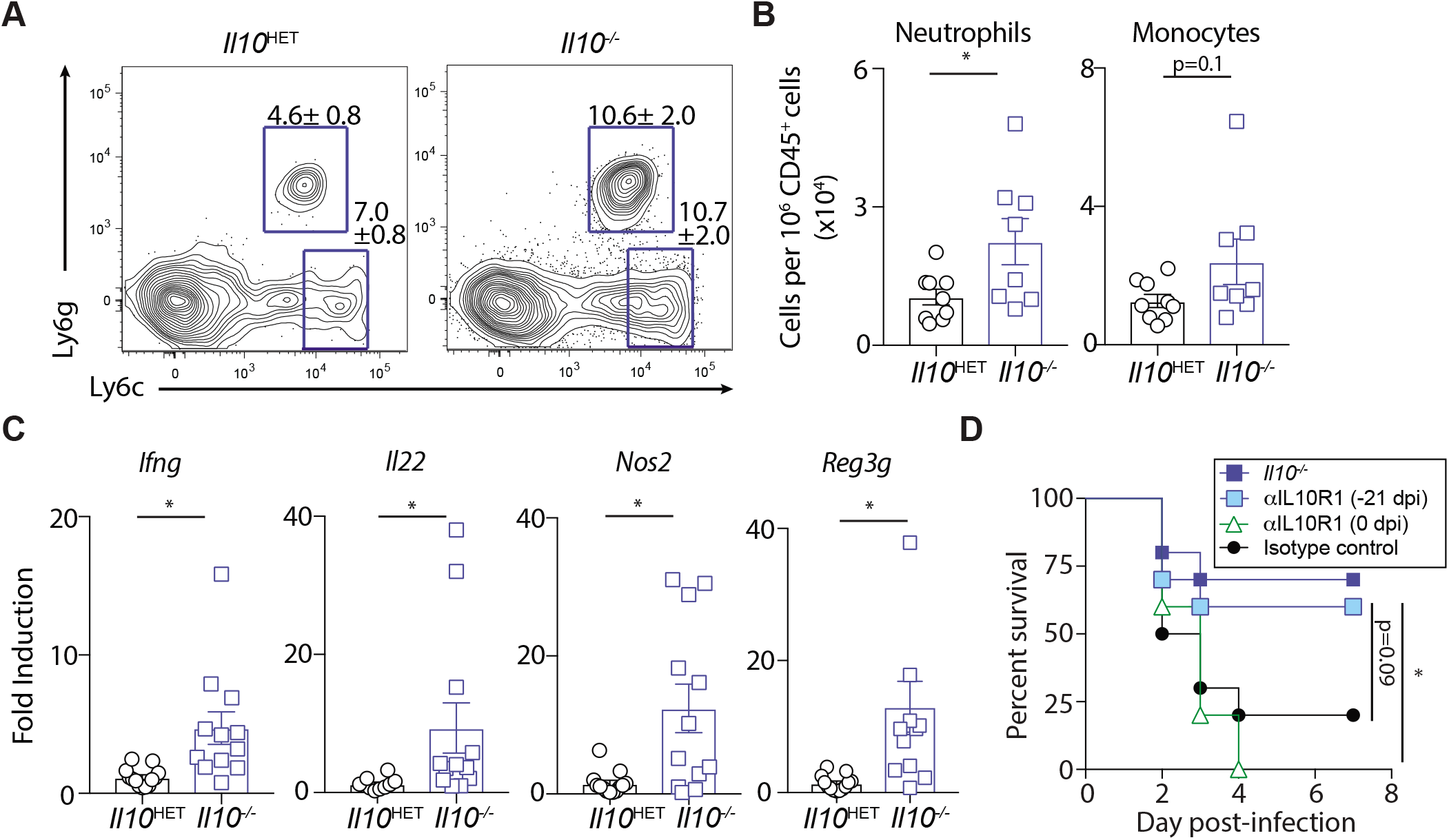
Loss of IL-10 signaling enhances intestinal immune activation prior to infection and decreases susceptibility to acute *C. difficile* infection. Antibiotic-treated uninfected and *Il10*^-/-^ mice and *Il10*^HET^ mice were sacrificed at the day of infection (prior to inoculation). **(A)** Frequency of neutrophils and monocytes in the large intestine lamina propria. FACS plots gated on live, CD45^+^, Non-T, Non-B cells, Siglec-F^neg^, CD11b^+^ cells. **(B)** Total number of neutrophils and monocytes in the large intestine lamina propria. Data is a combination representative of two independent experiments. *Il10*^-/-^,n=8; *Il10*^HET^, n=9. **(C)** Fold induction of type-1 and type-17 associated effector molecules in the colon of antibiotic-treated, uninfected *Il10*^-/-^ mice relative to antibiotic treated uninfected *Il10*^HET^ mice and normalized to *Hprt*. Data is a combination representative of three independent experiments. *Il10*^-/-^,n=12; *Il10*^HET^, n=13. Data shown are mean ± SEM. **(D)** C57BL/6 mice were cohoused with *Il10^-/-^* mice for two weeks then were administered anti-IL10R1 or isotype control (Rat IgG1) by i.p. injection weekly for three weeks prior to infection or received a single dose of anti-IL10R1 on the day of *C. difficile* infection and assessed for survival following infection. Data are a combination of two independent experiments (n=8 per group). * = p < 0.05. Statistical significance was calculated by an unpaired t-test or a log-rank test.

To determine whether loss of IL-10 signaling prior to infection and subsequent immune activation augments protection following *C. difficile* infection, IL-10 signaling was selectively blocked in C57BL/6 mice starting either three weeks prior to infection or once on the day of infection and survival was assessed. Antibody-mediated blockade of the IL-10-specific receptor IL-10R1 (αIL10R1) administered once a week for at least 3 weeks abrogates IL-10 signaling and replicates the intestinal inflammation observed in germline *Il10^-/-^* mice^49^. C57BL/6 mice that received weekly αIL10R1 treatment starting 3 weeks prior to infection exhibited a comparable survival to *Il10^-/-^* mice and significantly improved survival compared to C57BL/6 mice administered αIL10R1 at day 0 p.i. (Fig. 4D). These data support the hypothesis that IL-10 inhibits basal activation of intestinal immune defense genes prior to infection thereby rendering the host more susceptible to *C. difficile* infection.

### IL-22 is critical for host defense against *C. difficile* infection in *Il10^-/-^* mice

Inhibition of IL-10 signaling limits *C. difficile* pathogenesis only if initiated weeks prior to infection (Fig. 4D). Further, *Il10* deficiency leads to enhanced colonic *il22* and *ifng* expression in uninfected mice (Fig 4C). Both IL-22 and IFN-γ are critical in mounting a protective innate immune response during acute *C. difficile* infection^13,14,50^. These observations suggest immune activation prior to infection promotes improved survival in *Il10^-/-^* mice. To determine the relative contribution of these cytokine pathways to host protection in *Il10^-/-^* mice, *Il22* or *Tbx21* (the gene that encodes T-bet, a master transcription factor that regulates IFN-γ production) was genetically ablated in *Il10^-/-^* mice. Following *C. difficile* infection, *Il10.Tbx21* double knockout (dKO) mice exhibited comparable survival to *Il10^-/-^* mice, suggesting the IFN-γ pathway was dispensable for protection in *Il10^-/-^* mice (Fig. 5A). In contrast, *Il10.Il22* dKO mice were acutely susceptible to *C. difficile* infection (Fig. 5A). To confirm the dependence of IL-22 signaling for host protection in an IL-10 deficient setting, cohoused C57BL/6 or *Il10*^HET^ mice, *Il10^-/-^, Il22^-/-^, Il10.Il22* dKO, and *Il10r2*^-/-^ mice (IL-10R2 is the shared receptor subunit necessary for both IL-10 and IL-22 signaling) were pre-treated with antibiotics and infected with *C. difficile*. Genetic ablation of IL-22 signaling in an IL-10-deficient setting (*Il10.Il22* dKO and *Il10r2^-/-^* mice) led to significantly increased disease morbidity at day 2 p.i. (Fig. 5B), and mortality compared to *Il10^-/-^* mice (Fig. 5C). Collectively, these data support the conclusion that loss of IL-10 signaling leads to activation of IL-22 dependent host defense mechanisms that limit *C. difficile* pathogenesis.

**Figure 5.**
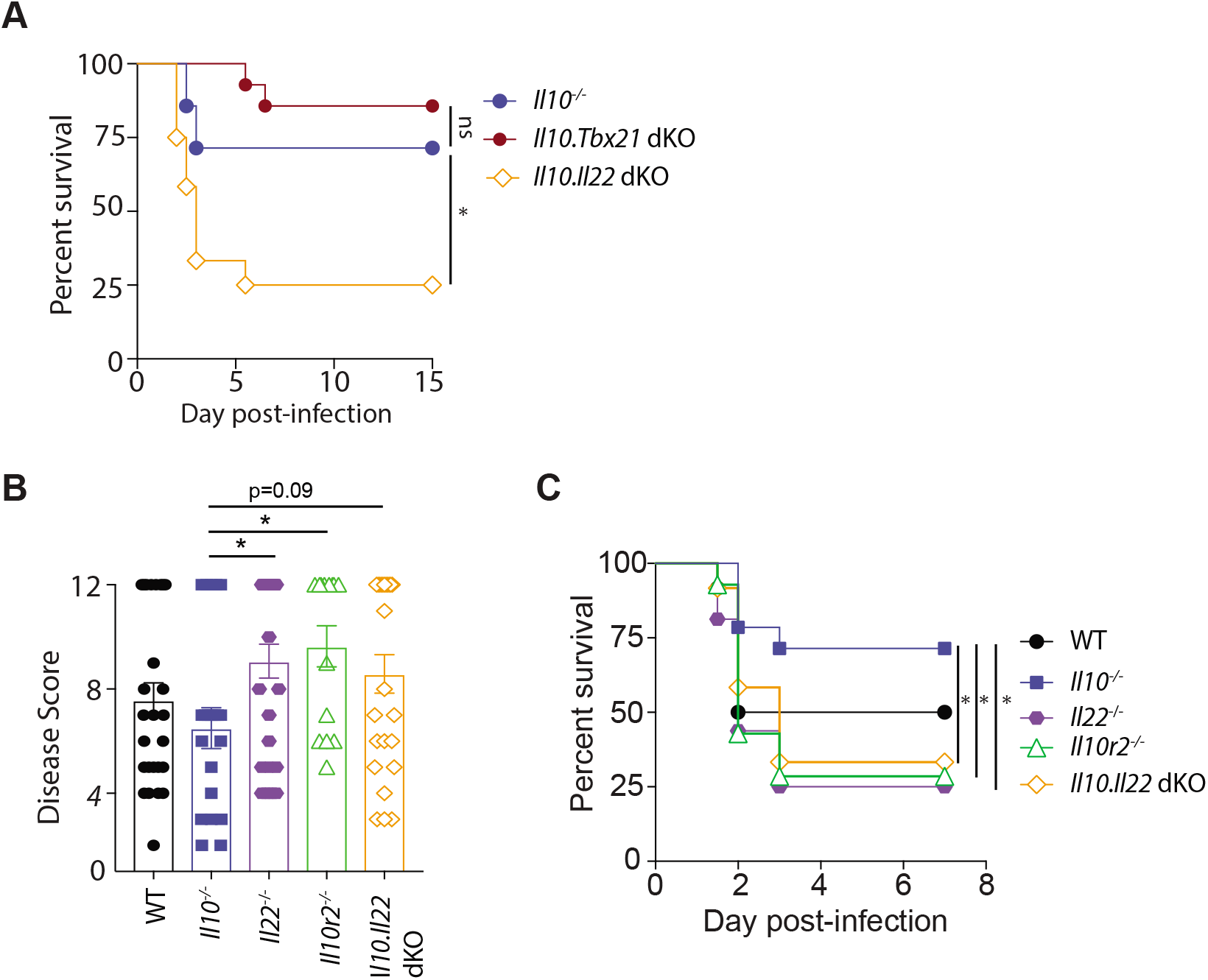
IL-22 signaling is required for protection against *C. difficile* infection in *Il10^-/-^* mice. **(A)** Cohoused *Il10^-/-^, Il10.Il22* dKO and *Il10.Tbx21* dKO mice were pre-treated with antibiotics and inoculated with approximately 400 spores of *C. difficile* (VPI 10463 strain) and assessed for survival following infection. Survival curve is a combination of three independent experiments. (*Il10^-/-^*, n=7; *Il10.Il22* dKO, n=12; *Il10.Tbx21* dKO, n=14). **(B)** Disease severity at day 2 p.i. and **(C)** survival curve of cohoused C57BL/6 or *Il10*^HET^ (wild-type - WT), *Il10^-/-^, Il10r2^-/-^, Il10^-/-^, Il22^-/-^*, and *Il10.Il22* dKO mice following *C. difficile* infection. Data shown are a combination of four independent experiments (WT, n=12; *Il10^-/-^*,n=14; *Il10r2^-/-^,n=14 Il22^-/-^*, n=16; *Il10.Il22* dKO, n=12). * = p<0.05. Statistical significance was calculated by an unpaired t-test or a log-rank test.

## Discussion

*C. difficile* infection induces a robust innate inflammatory response that has been extensively studied in the context of pathogenesis. Here, we employed *Il10^-/-^* mice to decouple constitutive intestinal inflammation from *C. difficile* infection-induced inflammation and determine their respective roles in pathogen defense. Collectively, our data support the conclusion that the absence of IL-10 signaling elevates host defenses prior to infection leading to reduced *C. difficile* pathogenesis in an IL-22 dependent manner. These results implicate a novel and deleterious role for IL-10 in dampening the IL-22 response during enteric *C. difficile* infection.

Previous work by Kim *et al.* assessed IL-10 in the context of *C. difficile* infection and observed severe histopathology in *C. difficile* infected *Il10^-/-^* mice infected at day 7 p.i. compared to cohoused C57BL/6 mice that developed mild *C. difficile* disease (10-15% weight loss; 100% survival in wild-type mice)^51^. The data in this report complements this finding and investigates the role of IL-10 in the acute response to severe *C. difficile* infection (20-30% weight loss; 25-50% survival in C57BL/6 mice). Together, these studies support a model where basal immune activation prior to infection can limit severe acute pathogenesis. However, prolonged *Il10* deficiency during a milder form of disease tips the balance toward inflammation-driven tissue immunopathology.

Despite the protective capacity of intestinal inflammation reported in this study in the context of *Il10* deficiency, expression of proinflammatory molecules does not uniformly limit *C. difficile* disease. Notably, *Il10^-/-^* mice are also a widely used model of inflammatory bowel disease (IBD), a well-appreciated risk factor for *C. difficile* disease^8,52^. Due to its multifactorial nature, however, clinical reports that link IBD to *C. difficile* are not able to differentiate what features of IBD drive increased *C. difficile* disease severity^52^. Research investigating the connection between IBD and *C. difficile* has employed mice treated with dextran sodium sulfate (DSS), a model of chemically-induced colitis, to disentangle the role of proinflammatory immune components in pathogenesis. DSS-treated mice exhibit increased susceptibility to *C. difficile* infection.^53^ and Saleh *et al.* demonstrated that IL-17 competent CD4^+^ T_H_17 cells activated by DSS colitis were sufficient to increase susceptibility to *C. difficile* infection in non-DSS treated mice^16^. This work identifies induction of IL-17 during IBD as an inflammatory pathway that promotes *C. difficile* disease.

In the context of the *C. difficile* infection, the IL-23/IL-22/IL-17 axis has a nuanced role in pathogenesis. Genetic ablation of IL-23, a cytokine upstream of IL-17, protects mice from severe *C. difficile* infection^17^ while mice deficient in IL-22, which is also directly downstream of IL-23, are acutely susceptible to infection^13,54^. Further, wild-type mice that receive rIL-22 treatment prior to *C. difficile* challenge are protected from severe infection^13^. Altogether, these observations, along with the results presented here, support a protective role for elevated IL-22 production. At the same time, induction of the IL-23 proinflammatory axis in the absence of IL-22, or in favor of IL-17 production, could drive more severe disease. Multiple IL-22 dependent mechanisms that mediate protection against *C. difficile* infection have been described. Hasegawa *et al.* demonstrated the induction of complement proteins via IL-22 signaling on hepatocytes is required to limit non-*C. difficile* bacterial translocation during severe *C. difficile* infection^13^. In addition to this systemic role for IL-22, a more recent study demonstrated a role for IL-22 signaling in modulation of intestinal epithelial glycosylation to enable growth of bacterial consumers of succinate, a crucial metabolite for *C. difficile* growth^55^. IL-22 also acts on intestinal epithelial cells to induce expression of genes that encode antimicrobial peptides, including RegIIIγ, lipocalin-2, and calprotectin^56–58^, that limit damage of adherent or mucosal-associated commensal bacteria to the epithelium^56,57^ the latter of which has been associated with host protection against *C. difficile*^59^. In support of this, Gunesekera *et al.* showed that *Il10^-/-^* mice exhibited enriched expression of these same IL-22 dependent antimicrobial genes, all of which were uniquely lost in *Il10.Il22* dKO mice^60^. Thus, elevated IL-22 expression at the time of *C. difficile* infection, as observed in *Il10*^-/-^ mice, positions the host to limit toxin-mediated destruction of the epithelial barrier.

Immune activation in IL-10 deficient hosts disrupts homeostasis at steady-state but is also beneficial in the context of an acute *C. difficile* infection by elevating baseline defense mechanisms prior to infection. This concept of immunological tuning prior to infection has been previously observed with commensal bacteria providing tonic signaling to maintain antiviral defenses in a poised state of readiness to rapidly respond upon viral infection^61–63^. Thus, the trade-off of constitutive intestinal IL-10 expression is diminished basal activation of immune defense genes and therefore a decreased capacity of the host to respond to pathogen challenge. Understanding the dynamics of this biological balancing act could help develop therapies that selective or transiently targeting the protective components of immune activation in at-risk patients while avoiding deleterious side effects of prolonged inflammation.

## Materials and Methods

### Mice

Four to six-week old wild-type C57BL/6J, *Il10^-/-^, Tbx21^-/-^, Il10rb^-/-^*mice were purchased from the Jackson Laboratory. *Il22^-/-^* mice were provided by R. Flavell (Yale University). All knockout mouse strains were derived on a C57BL/6 background. All mice were bred and maintained in autoclaved cages under specific pathogen-free conditions at the University of Pennsylvania. All experiments with cohoused C57BL/6 and *Il10^-/-^* mice were done at Memorial Sloan Kettering Cancer Center. *Il10Il22* dKO and *Il10.Tbx21* dKO mice were generated by breeding *Il10^-/-^* mice with *Il22^-/-^* and *Tbx21^-/-^* mice, respectively. The presence of *Helicobacter spp*., a bacterial genus sufficient to trigger intestinal inflammation in *Il10^-/-^* mice by three months of age^64^, was confirmed by PCR in the feces of all breeder *Il10^-/-^* mice and *Il10*^-/-^-derived mouse strains. Sex- and age-matched control mice were used in all experiments according to institutional guidelines for animal care. All animal procedures were approved by the Institutional Animal Care and Use Committee of the University of Pennsylvania and Memorial Sloan Kettering Cancer Center.

### Antibiotic Pretreatment, *C. difficile* infection, and Mouse Monitoring

Mice were cohoused for 3 weeks prior to antibiotic treatment and then supplemented with metronidazole (0.25 g/l), neomycin (Sigma) (0.25 g/l), and vancomycin (Novaplus) (0.25 g/l) in drinking water for 3 days. One day following cessation of antibiotic water, mice received 200 mg of clindamycin (Sigma) by i.p. injection. Twenty-four hours later, mice received approximately 400 *C. difficile* spores (VPI10463 strain ATCC #43255) via oral gavage. For antibody-mediated blockade experiments, mice received 1 mg of αIL10R1 antibody (clone 1B1.3A, Bio X Cell) or mouse IgG1 isotype control (clone MOPC-21, Bio X Cell) i.p. weekly starting either 3 weeks prior to infection or at day of infection. After infection, mice were monitored and scored for disease severity by four parameters: weight loss (> 95% of initial weight = 0, 95%– 90% initial weight = 1, 90%–80% initial weight = 2, < 80% = 3), surface body temperature (> 32°C = 0, 32°C–30.5°C = 1,30.5°C–29°C = 2, < 29°C = 3), diarrhea severity (formed pellets = 0, loose pellets = 1, liquid discharge = 2, no pellets/caked to fur = 3), morbidity (score of 1 for each symptoms with a maximum score of 3; ruffled fur, hunched back, lethargy, ocular discharge). Mice that exhibited severe disease, defined as a surface body temperature below 29.5°C or weight loss in excess of 30%, were humanely euthanized.

### *C. difficile* Quantification

Fecal pellets or cecal content were resuspended in deoxygenated PBS, and ten-fold serial dilutions were plated anaerobically at 37° C on brain-heart infusion agar supplemented with yeast extract, L-cysteine, D-cycloserine, cefoxitin, and taurocholic acid (CCBHIS-TA) and colony-forming units (CFUs) were enumerated 24 hours later. Prior to infection, fecal samples from mice were cultured overnight in CCBHIS-TA liquid broth, then serially diluted and grown for 24 hours on CCBHIS-TA plates to ensure that mice did not harbor endogenous *C. difficile* in their microbiota. Supernatants from the cecal or fecal content were obtained after centrifugation for cytotoxicity assays and LCN-2 ELISA (Bethyl Labs).

### *C. difficile* Toxin Cytotoxicity Assay

Vero cells were seeded in 96-well plates at 1 x 10^4^ cells/well and incubated for 24 hours at 37°C in 5% CO_2_. Cecal or fecal supernatants were added in ten-fold dilutions to the Vero cells (100 μL/well) and incubated overnight prior to removal, rinsing with PBS, and replacement with fresh media. The presence of *C. difficile* toxins A and B was confirmed by neutralization with antitoxin antisera (Techlab, Blacksburg, VA). The data are expressed as the log10 reciprocal value of the last dilution where cell rounding was observed. Cell morphological changes were observed after 18 hours using a Nikon inverted microscope. The cytopathic effect was determined as rounded cells in comparison to the negative control wells.

### 16S rRNA sequencing

Cecal content was collected from uninfected mice and mice infected with *C.difficile* at 2 dpi. DNA was extracted using the Qiagen MagAttract Power Microbiome kit DNA/RNA kit (Qiagen, catalog no. 27500-4-EP) and used for rRNA sequencing and *Helicobacter spp.* PCR. Genus-specific PCR was conducted on purified bacterial DNA from feces of *Il10^-/-^* breeder mice and The V4-V5 region of the 16S rRNA gene was amplified from each sample using the dual indexing sequencing strategy as described previously^65^. Sequencing was done on the Illumina MiSeq platform. The V4-V5 region of the 16S rRNA gene was sequenced and demultiplexed using the fqgrep tool. Data were imported into QIIME2 (v. 2020.2)^66^ and denoised using the DADA2 plugin^67^. For data in Supplemental Figure 3, the fqgrep tool (https://github.com/indraniel/fqgrep) was used to demultiplex the sequences followed by denoising using the DADA2 (v. 1.14.1)^67^ implementation in R (v. 3.6.3)^68^. Due to quality issues on the reverse reads, only the forward reads were used for denoising. Datasets were taxonomically classified in QIIME2 using the q2-feature-classifier^69^ classify-sklearn naïve Bayes classifier with a newly generated classifier against Greengenes 13_8 99% OTU sequences^70^. Phylogenetic trees were generated using mafft^71^ and the q2-phylogeny plugin^72^. Data were then imported into R for further analyses with phyloseq (v. 1.30.0)^73^ and visualization with ggplot2 (v. 3.3.0)^74^. Unweighted UniFrac^75^ dissimilarity was calculated to generate PCoA plots and for creating dendrograms using the hclust function (stats package in R core, v. 3.6.3). Finally, a linear model was built using the lm() and padjust() functions (stats package, v. 3.6.3) as well as the tidyverse package (v. 1.3.0)^76^.

### Isolation of lamina propria cells and flow cytometry

Single cell suspensions were obtained from the large intestine lamina propria compartment by longitudinally cutting the large intestine and washing out its contents in PBS. Intestinal tissues were incubated at 37°C under gentle agitation in stripping buffer (PBS, 5 mM EDTA, 1 mM dithiothreitol, 4% FCS, 10μg/mL penicillin/streptomycin) for 10 minutes to remove epithelial cells and then for another 20 minutes for the IEL. The tissue was digested with collagenase IV 1.5 mg/mL (500 U/mL) and DNase (20 μg/mL) in complete media (DMEM supplemented w/ 10% FBS, 10 μg/mL penicillin/streptomycin, 50 μg/mL gentamicin, 10 mM HEPES, 0.5 mM β-mercaptoethanol, 20 μg/mL L-glutamine) for 30 minutes at 37°C under gentle agitation. Supernatants containing the lamina propria fraction were passed through a 100 μm then subsequently 40 μm cell strainer. After counting, cells were plated at 10^6^ cells per well in 96-well round bottom plates and washed twice in 1x PBS before incubating with a cell viability dye 20 minutes at room temperature (Invitrogen AQUA dye). After Fc blockade (anti-mouse CD16/32 TruStain, BioLegend), cells were stained using a standard flow cytometric staining protocol with fluorescently conjugated antibodies specific to CD3ε, CD5, CD19, CD45, MHC-II, Siglec-F, Ly-6G, Ly6C, CD11b, and CD11c. Stained cells were kept in FACS buffer at 4°C until run. Samples were acquired on an LSR-II flow cytometer (Becton Dickinson). Data were analyzed using FlowJo version 9.9.6 software. Cell populations were calculated from total cells per colon as a percentage of live CD45^+^ cells.

### Cytokine and Chemokine Quantification

Cecal tissue was homogenized in tissue extraction buffer with protease inhibitors for 1 minute by bead beating with steel beads. Homogenates were centrifuged at x10,000 rpm for 5 minutes and supernatants were collected and stored at −80° C. Supernatants containing protein were analyzed by Mouse Multiplex Luminex assay (Invitrogen) at the Human Immunology Core at University of Pennsylvania. Concentrations displayed as ng of analyte per gram of cecal tissue.

### Tissue RNA Isolation, cDNA Preparation, and qRT-PCR

RNA was isolated from proximal colon tissue using mechanical homogenization and Trizol isolation (Invitrogen) according to the manufacturer’s instructions. cDNA was generated using QuantiTect reverse transcriptase (QIAGEN). Quantitative RT-PCR was performed on cDNA using either TaqMan primers and probes or QuantiTect primers in combination with TaqMan PCR Master Mix (ABI) or SYBR Green chemistry and reactions were run on a RT-PCR system (QuantiStudio 6 Flex, Applied Biosystems). Gene expression is displayed as fold increase over antibiotic-pre-treated, uninfected control mice and was normalized to the *Hprt* gene.

### Statistical Analysis

Results represent means ± SEM. Statistical significance was determined by the unpaired t-test and log-rank test for survival curve. Statistical analyses were performed using Prism GraphPad software v6.0 (* p< 0.05; ** p< 0.01; *** p < 0.001).

---

**Supplementary Figure 1: *Il10^-/-^* mice are less susceptible to *C. difficile* infection compared to cohoused C57BL/6 mice.** (**A)** *Il10^-/-^* and C57BL/6 (B6) mice were inoculated with approximately 400 spores of *C. difficile* (VPI 10463 strain) and assessed for survival following infection. Survival curve is a combination of four independent experiments (*Il10*^-/-^, n=31; C57BL/6, n=33). **(B)** *C. difficile* burden and **(C)** toxin levels in the cecal content at days 2 and 4 p.i. ** = p<0.01. Statistical significance was calculated by a log-rank test.

**Supplementary Figure 2: Identification of individual ASVs that correlate with *C. difficile-infected Il10^-/-^* mice and *Il10*^HET^ mice. (A)** Linear modeling of ASV abundances in IL10^HET^ and IL10^-/-^ cecal microbiotas fail to identify differences in microbiota compositions. **(A)** Abundance of top 3 ASVs correlating significantly with experimental groups were plotted. **(B)** Model comparisons for experimental groups show significantly different ASV abundances that correlate with group phenotype. *p < 0.05, **p < 0.01, ***p <0.001.

**Supplementary Figure 3: Cohousing *Il10^-/-^* mice with C57BL/6 mice assimilates their microbiota prior to infection.** Fecal pellets were collected from *Il10^-/-^* and wild-type mice starting prior to cohousing (day −64 p.i.), following cohousing (day −55, −47 p.i.), the start of antibiotic treatment (day −6 p.i.), and the day of infection (day 0 p.i.). Fecal pellets were processed for 16S rRNA bacterial gene profiling. **(A)** Unweighted UniFrac principal coordinate analysis plot of 16S bacterial rRNA ASVs. **(B)** Relative abundance of top 15 bacterial ASVs. **(C)** Dendrogram representation of intestinal microbial communities using unsupervised hierarchical clustering of unweighted UniFrac distances to identify similarities between samples. **(D)** Microbial alpha diversity as determined by the Shannon diversity index. **(E)** LEfSe analysis identifying significantly differentially abundance ASVs prior to cohousing (day −64 p.i.), prior to antibiotics (day −6 p.i.), and at the day of infection (day 0 p.i.). ** = p<0.01.

**Supplementary Figure 4: *Il10^-/-^* and *Il10*^HET^ mice exhibit comparable granulocyte infiltration induction of proinflammatory cytokines and chemokines following acute *C. difficile* infection.** *Il10^-/-^* and *Il10*^HET^ mice were inoculated with approximately 400 spores of *C. difficile* (VPI 10463 strain) or mock infected and sacrificed two days later. **(A-B)** Flow cytometry gating strategy identifying frequency of **(A)** eosinophils, **(B)** neutrophils, monocytes in the large intestine lamina propria of day 2 p.i. *Il10^-/-^* and *Il10*^HET^ mice. First FACS plot is gated on live, CD45^+^ cells. **(C)** Proinflammatory cytokines and **(D)** chemokines protein levels in the cecal tissue homogenate. Data shown are a combination of two independent experiments (uninfected *Il10*^-/-^,n=7; uninfected *Il10*^HET^, n=6; day 2 infected *Il10*^-/-^,n=8; uninfected *Il10*^HET^, n=7). Data shown are mean ± SEM.

**Supplemental Table 1.** PERMANOVA analysis of unweighted UniFrac distances between the intestinal microbial communities of antibiotic-treated uninfected and *C. difficile* infected *Il10^-/-^* and *Il10*^HET^ mice at day 2 p.i. Statistical tests performed on data displayed in Figure 2.

**Supplemental Table 2.** PERMANOVA analysis of unweighted UniFrac distances between the intestinal microbial communities of *Il10^-/-^* and C57BL/6 mice starting prior to cohousing (day −64 p.i.), following cohousing (day −55, −47 p.i.), the start of antibiotic treatment (day −6 p.i.), and the day of infection (day 0 p.i.). Statistical tests performed on data displayed in Supplemental Figure 3.

## Acknowledgements

We thank the members of the Abt lab for helpful discussion and critical reading of the manuscript. We would also like to thank L. Mattei of the Penn CHOP Microbiome Core and L. Lang of the Lucille Castori Center for Microbes, Inflammation and Cancer for technical expertise in high throughput sequencing and E. Pamer for mice strains. Finally, we thank L. Zhao and R. Shimol of the Penn Human Immunology Core for technical expertise with Luminex assays. This research was supported by the NIH (R00 AI125786 to M.C.A and T32 AI141393 to E.S.C.).

