## Supplemental Figures for "Loss of IL-10 signaling promotes IL-22 dependent host defenses against acute *Clostridioides difficile* infection"

Supplemental Figure 1

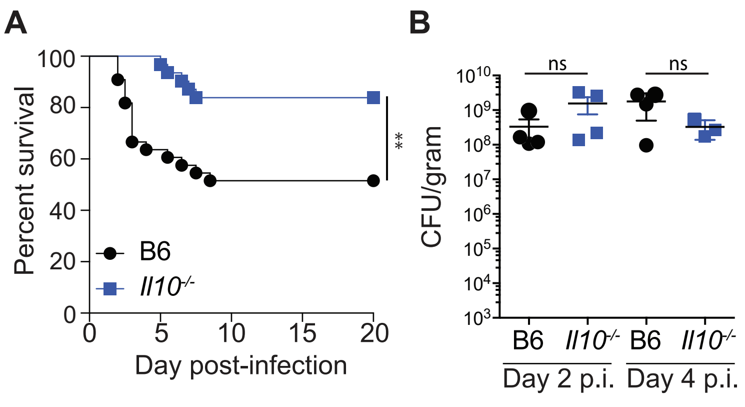


**Supplementary Figure 1: Il10-/- mice are less susceptible to C. difficile infection compared to cohoused C57BL/6 mice.** (A) Il10-/- and wild-type mice were inoculated with approximately 400 spores of C. difficile (VPI 10463 strain) and assessed for survival following infection. Survival curve is a combination of four independent experiments (Il10-/-, n=31; C57BL/6, n=33). (B) C. difficile burden and (C) toxin levels in the cecal content at days 2 and 4 p.i. ** = p<0.01. Statistical significance was calculated by a log-rank test.

Supplemental Figure 2

**A**

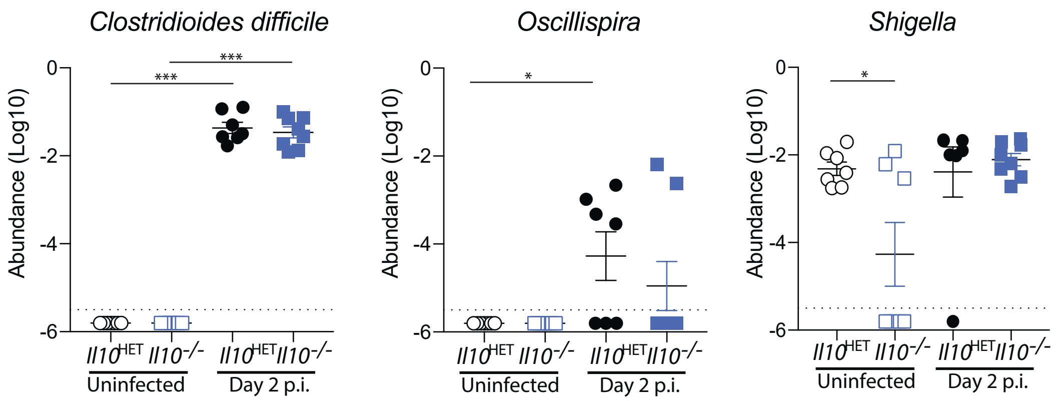


**B**

| Linear Regression Model | ASV | p-value | FDR | Significance |
|---|---|---|---|---|
| *Il10*$^{HET}$ uninf vs *Il10*$^{-/-}$ uninf | *Shigella* | 0.022 | 0.044 | * |
| | *Lactobacillus vaginalis* | 0.049 | 0.096 | ns |
| *Il10*$^{HET}$ uninf vs *Il10*$^{HET}$ Day 2 | *Clostridioides difficile* | $1.01 \times 10^{-13}$ | $3.15 \times 10^{-13}$ | *** |
| | *Oscillospira* | 0.017 | 0.033 | * |
| | *Aggregatibacter pneumotropica* | 0.028 | 0.056 | ns |
| *Il10*$^{-/-}$ uninf vs *Il10*$^{-/-}$ Day 2 | *Clostridioides difficile* | $1.39 \times 10^{-14}$ | $4.67 \times 10^{-14}$ | *** |
| *Il10*$^{HET}$ Day 2 vs *Il10*$^{-/-}$ Day 2 | No ASVs identified | n/a | n/a | n/a |

**Supplementary Figure 2: Identification of individual ASVs that correlate with C. difficile-infected Il10-/- mice and Il10HET mice.** (A) Linear modeling of ASV abundances in IL10HET and IL10-/- cecal microbiotas fail to identify differences in microbiota compositions. (A) Abundance of top 3 ASVs correlating significantly with experimental groups were plotted. (B) Model comparisons for experimental groups show significantly differ-ent ASV abundances that correlate with group phenotype. *p < 0.05, **p < 0.01, ***p <0.001.

Supplemental Figure 3

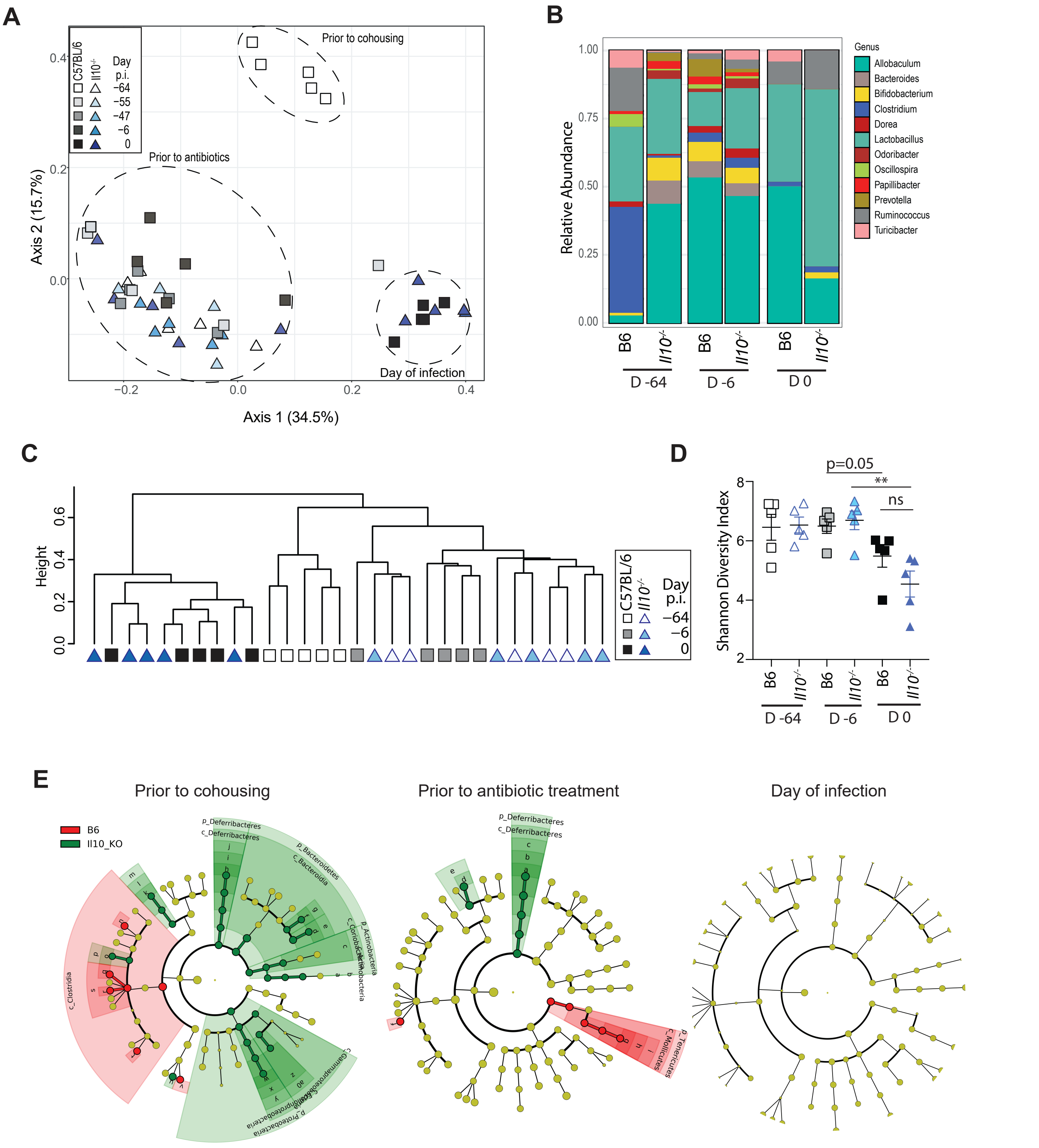


**Supplementary Figure 3: Cohousing Il10-/- mice with C57BL/6 mice assimilates their microbiome prior to infection.** Fecal pellets were collected from Il10-/- and wild-type mice starting prior to cohousing (day -64 p.i.), following cohousing (day -55, -47 p.i.), the start of ABX treatment (day -6 p.i.), and the day of infection (day 0 p.i.). Fecal pellets were processed for 16S rRNA bacterial gene profiling. (A) Unweighted UniFrac principal coordinate analysis plot of 16S bacterial rRNA ASVs. (B) Relative abundance of top 15 bacterial ASVs. (C) Dendrogram representation of intestinal microbial communities using unsupervised hierarchical clustering of unweighted UniFrac distances to identify similarities between samples. (D) Microbial alpha diversity as determined by the Shannon diversity index. (E) LEfSe analysis identifying significantly differentially abundance ASVs prior to cohousing (day -64 p.i.), prior to ABX (day -6 p.i.), and at the day of infection (day 0 p.i.). ** = p<0.01.

Supplemental Figure 4

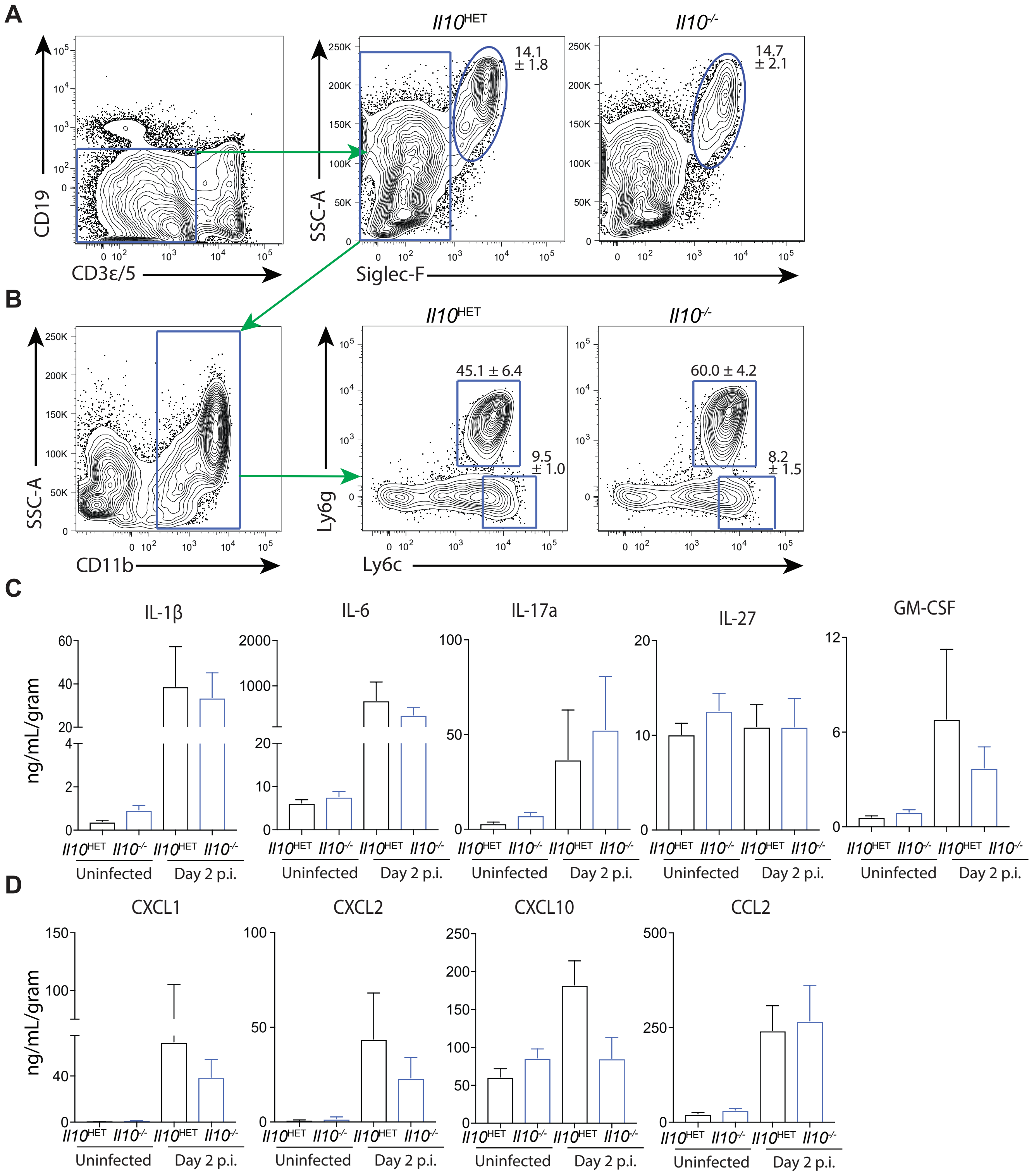


**Supplementary Figure 4: Il10-/- and Il10HET mice exhibit comparable granulocyte infiltration induction of proinflammatory cytokines and chemokines following acute C. difficile infection.** Il10-/- and Il10HET mice were inoculated with approximately 400 spores of C. difficile (VPI 10463 strain) or mock infected and sacrificed two days later. (A-B) Flow cytometry gating strategy identifying frequency of (A) eosinophils, (B) neutrophils, monocytes in the large intestine lamina propria of day 2 p.i. Il10-/- and Il10HET mice. First FACS plot is gated on live, CD45+ cells. (C) Proinflammatory cytokines and (D) chemokines protein levels in the cecal tissue homogenate. Data shown are a combination of two independent experiments (uninfected Il10-/-,n=7; uninfected Il10HET, n=6; day 2 infected Il10-/-,n=8; uninfected Il10HET, n=7). Data shown are mean ± SEM.

Supplemental Table 1

| Pair Comparison | $R^2$ | p-value | FDR | Significance |
|---|---|---|---|---|
| *Il10*$^{HET}$ uninf vs *Il10*$^{-/-}$ uninf | 0.0724 | 0.542 | 0.650 | ns |
| *Il10*$^{HET}$ Day 2 vs *Il10*$^{-/-}$ Day 2 | 0.0457 | 0.932 | 0.932 | ns |
| *Il10*$^{HET}$ uninf vs *Il10*$^{HET}$ Day 2 | 0.1005 | 0.140 | 0.420 | ns |
| *Il10*$^{-/-}$ uninf vs *Il10*$^{-/-}$ Day 2 | 0.0712 | 0.410 | 0.650 | ns |

**Supplemental Table 1.** PERMANOVA analysis of unweighted UniFrac distances between the intestinal microbial communities of ABX-treated uninfected and C. difficile infected Il10-/- and Il10HET mice at day 2 p.i. Statistical tests performed on data displayed in Figure 2.

Supplemental Table 2

| Pair Comparison | $R^2$ | p-value | FDR | Significance |
|---|---|---|---|---|
| *Il10*$^{-/-}$ D -64 vs C57BL/6 D -64 | 0.5607 | 0.009 | 0.021 | * |
| *Il10*$^{-/-}$ D -55 vs C57BL/6 D -55 | 0.1123 | 0.383 | 0.421 | ns |
| *Il10*$^{-/-}$ D -47 vs C57BL/6 D -47 | 0.1831 | 0.010 | 0.021 | * |
| *Il10*$^{-/-}$ D -6 vs C57BL/6 D -6 | 0.1714 | 0.099 | 0.126 | ns |
| *Il10*$^{-/-}$ D 0 vs C57BL/6 D 0 | 0.1274 | 0.303 | 0.351 | ns |

**Supplemental Table 2.** PERMANOVA analysis of unweighted UniFrac distances between the intestinal microbial communities of Il10-/- and C57BL/6 mice starting prior to cohousing (day -64 p.i.), following cohousing (day -55, -47 p.i.), the start of ABX treatment (day -6 p.i.), and the day of infection (day 0 p.i.). Statistical tests performed on data displayed in Supplemental Figure 3.
